# Enterovirus replication and dissemination are differentially controlled by type I and III interferons in the GI tract

**DOI:** 10.1101/2022.02.07.479406

**Authors:** Alexandra I. Wells, Kalena A. Grimes, Carolyn B. Coyne

## Abstract

Enteroviruses are amongst the most common viral infectious agents of humans and cause a broad spectrum of mild-to-severe illness. Enteroviruses are primarily transmitted by the fecal-oral route, but the events associated with their intestinal replication *in vivo* are poorly defined. Here, we developed a neonatal mouse model of enterovirus infection by the enteral route using echovirus 5 and used this model to define the differential roles of type I and III interferons (IFNs) in enterovirus replication in the intestinal epithelium and subsequent dissemination to secondary tissues. We show that human FcRn, the primary receptor for echoviruses, is essential for intestinal infection by the enteral route and that type I IFNs control dissemination to secondary sites, including the liver. In contrast, type III IFNs limit enterovirus infection in the intestinal epithelium and mice lacking this pathway exhibit persistent epithelial replication. Finally, we show that echovirus infection in the small intestine is cell-type specific and occurs exclusively in enterocytes. These studies define the type-specific roles of IFNs in enterovirus infection of the GI tract and the cellular tropism of echovirus intestinal replication.

## Introduction

Enteroviruses are small (∼30nm) single stranded RNA viruses that are comprised of coxsackieviruses (CVA and CVB), rhinoviruses, poliovirus (PV), enteroviruses 71 and D68 (e.g., EV-71, EV-D68), and echoviruses (which includes ∼30 serotypes). Echoviruses can cause 15-30% of nosocomial infections is Neonatal Intensive Care Units (NICUs) and often result in aseptic meningitis and liver failure, which can be fatal^1–4^. The National Enterovirus Surveillance System (NESS) indicates that between the years of 2014-2016 echoviruses were amongst the most commonly circulating enteroviruses in the U.S.^5^. Globally, outbreaks of other echoviruses including echovirus 5 (E5) have been associated with a range of clinical outcomes, with most severe disease occurring in infants and children^6–8^.

Enteroviruses are primarily transmitted through the fecal-oral route and initiate host entry via the epithelial lining of the GI tract. We have previously shown that echoviruses robustly infect human stem cell-derived intestinal enteroids and exhibit a cell-type specificity of infection, with preferential infection in enterocytes and enteroendocrine cells^9,10^. Additionally, echovirus infections cause damage to barrier function in enteroid-derived intestinal epithelial monolayer cultures^10^, suggesting that virus-mediated epithelial damage could contribute to dissemination from the intestine. The impact of host innate immune signaling on enteroviral infections in the intestinal epithelium is largely unknown. Previous studies in mouse models using PV and EV71 have shown that ablation of type I interferon (IFN) signaling by deletion of the IFNα/β receptor (IFNAR) is required for infection by the oral route^11–13^, suggesting that these IFNs play a central role in the protection of the GI tract from enterovirus infection. Whether this enhancement was the result of increased infection in the intestinal epithelium directly and/or resulted from alterations in infection of non-epithelial cell types remains unclear. Other studies using CVB show that infection by the enteral route remains inefficient in IFNAR^-/-^ animals, suggesting that these IFNs are not involved in intestinal innate immune responses^14^. Consistent with this, type III IFNs, which are comprised of IFN-λs 1-3 in humans, are preferentially induced in enterovirus-infected human enteroids^9,10^. For other enteric viruses, such as reoviruses, rotaviruses, and noroviruses, type III IFNs specifically control viral replication in the intestinal epithelium *in vivo*, with type I IFNs impacting the lamina propria^11,15–18^. Thus, the roles of type I and III IFNs in the control of enteroviral infections *in vivo* remain unclear.

We and others previously identified the human neonatal Fc receptor (hFcRn) as a primary receptor for echoviruses^19,20^. In contrast, mouse FcRn does not function as an echovirus receptor and does not support replication *in vivo*^19,21^. FcRn is expressed at the apical membrane of polarized enterocytes, where it binds to IgG and albumin, is internalized by endocytosis, and delivers its cargo to early and late endosomes, with the eventual release of IgG and albumin into the interstitium^22^. However, expression of hFcRn alone is not sufficient for echovirus infection in adult or neonatal mice and ablation of IFNAR in hFcRn-expressing mice is required for infection following intraperitoneal (IP) inoculation^21^. However, the roles of hFcRn and IFN signaling following inoculation via the enteral route was not explored.

Here, we established *in vitro* and *in vivo* models to define the impact of hFcRn expression and type I and III IFN signaling in echovirus infections in the GI tract. To do this, we generated mice expressing hFcRn that are deficient in the type III IFN receptor (IFNLR) and compared their susceptibility to enteral echovirus infection to hFcRn-expressing mice lacking IFNAR expression, or immunocompetent animals expressing hFcRn alone. Whereas expression of hFcRn was necessary and sufficient to support echovirus replication in primary murine stem cell-derived enteroids, it was not sufficient for infection of immunocompetent mice following oral gavage. We show that hFcRn-expressing mice deficient in IFNLR expression are unable to control echovirus infection in the GI tract and exhibit persistent replication in the intestinal epithelium, which occurred exclusively in enterocytes. However, these animals did not exhibit any morbidity or mortality from infection and there was no dissemination to secondary tissues. In contrast, there was robust dissemination in hFcRn-expressing mice deficient in IFNAR expression, which resulted in significant morbidity and mortality. However, we did not observe active replication in the intestinal epithelium of these animals. These findings define the differential roles of type I and III IFNs in the control of echovirus replication in the GI tract and in subsequent dissemination.

## Results

### Human FcRn is required for echovirus infection of murine-derived enteroids

To define the role of hFcRn in infections of the murine intestine, we generated neonatal enteroids from C57/BL6 (wild-type, WT) mice and mice expressing hFcRn (hFcRn^Tg32^). hFcRn^Tg32^ mice are deficient in expression of mouse FcRn and express human FcRn under the control of the native human promotor^23^. Stem cell-derived enteroids differentiated to form three-dimensional structures containing cells present in the epithelium *in vivo*, including enterocytes and mucin-secreting goblet cells (**Figure 1A**). Consistent with what has been described for murine fibroblasts derived from WT mice^19^, enteroids derived from WT mice were resistant to E5 infection (**Figure 1B**). In contrast, enteroids derived from hFcRn^Tg32^ mice were highly permissive to E5 infection, which peaked at ∼ 24 hours post inoculation (hpi) (**Figure 1B**). To define the host response of E5-infected enteroids, we performed bulk RNASeq followed by differential expression analysis. Similar to previous results in human enteroids^9^, murine-derived enteroids induced the selective expression of transcripts associated with the type III IFNs IFNλ-2 and IFNλ-3 (mice do not express IFN-λ1) (**Figure 1C**). Differential expression analysis revealed the induction of 48 transcripts in E5-infected hFcRn^Tg32^ enteroids, 42 of which are classified as interferon stimulated genes (ISGs) (**Figure 1D**), supporting a prominent role of IFN signaling in the intestinal innate immune response to echovirus infections.

**Figure 1.**
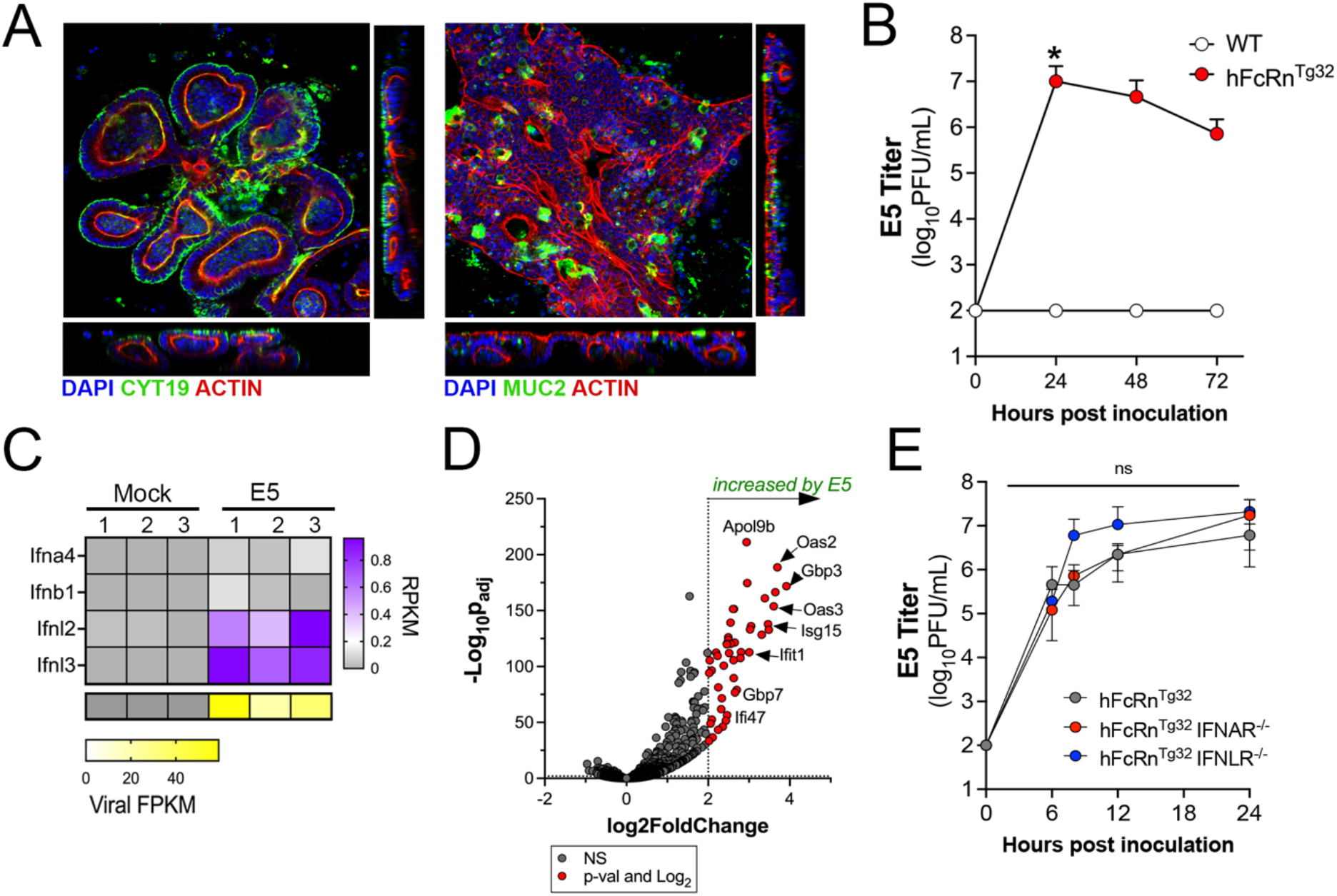
Human FcRn is necessary and sufficient for echovirus infection of murine-derived primary enteroids. **(A)** Murine enteroids were generated Lgr5^+^ crypts isolated from the small intestines of five 10-day old neonatal C57BL/6J (WT) mice. Confocal microscopy of enteroids immunostained with cytokeratin-19 in green and actin in red (left) or mucin-2 in green and actin in red (right) ∼10-days post-culturing. **(B)** WT (white) or hFcRn^Tg32^ (grey) enteroids were generated from small intestine tissue from 10-day old neonatal mice and infected with 10^6^ PFU of neutral red incorporated E5. Viral titers (log_10_PFU/mL) were assessed in cell culture supernatants at the indicated time points. **(C)** Heatmap of RPKM values of the type I IFNs Ifna4 and Ifnb1 and the type III IFNs Ifnl2 and Ifnl3 from bulk RNASeq of uninfected (mock)- or E5 infected hFcRn^Tg32^ enteroids at 24hrs post-infection. Key at right, purple indicates higher reads and grey denotes no reads detected. At bottom, viral FPKM values from samples shown at top. Yellow indicates high viral RNA reads and grey denotes no reads detected. **(D)** Volcano plot comparing differentially expressed transcripts in E5 infected hFcRn^Tg32^ enteroids compared to mock controls as determined by DeSeq2 analysis. Grey circles represent genes whose expression was not significantly changed. Red circles represent genes that were significantly changed by E5 infection. Significance was set at a p<0.01 and a log_2_fold-change of ± 2. **(E)** Enteroids generated from the small intestine of hFcRn^Tg32^ (grey), hFcRn^Tg32^-IFNAR^-/-^ (red), or hFcRn^Tg32^-IFNLR^-/-^ (blue) were infected with 10^6^ PFU of neutral red incorporated E5. Viral titers (log_10_PFU/mL) are shown at indicated time points. In (B) and (E), data are shown as mean ± standard deviation from three independent replicates. Enteroids were isolated from at least five 10-day old neonatal mice and were pooled together during the LGR5^+^ crypt isolation. Significance in B was determined by a Kruskal-Wallis test with a Dunn’s test for multiple comparisons. Significance in D was determined using a two-way ANOVA with a Geisser-Greenhouse correction and a Tukey’s multiple comparisons test. (*p<0.05, ns not significant.)

The selective induction of type III IFNs in murine-derived enteroids suggests that these IFNs are key mediators in the control of echovirus infections in the intestinal epithelium. To test this, we derived enteroids from small intestine tissue of mice expressing hFcRn that are deficient in IFNAR expression (hFcRn^Tg32^-IFNAR^-/-^)^21^. To perform parallel studies in enteroids deficient in type III IFN signaling, we crossed hFcRn^Tg32^ mice to mice deficient in IFNLR expression (hFcRn^Tg32^-IFNLR^-/-^). Enteroids were generated from the small intestines of immunocompetent hFcRn^Tg32^, hFcRn^Tg32^-IFNAR^-/-^, and hFcRn^Tg32^-IFNLR^-/-^ mice and the levels of E5 replication compared between these genotypes. We did not detect any significant differences in E5 replication between hFcRn-expressing immunocompetent enteroids and those deficient in either IFNAR or IFNLR expression (**Figure 1E**). These data show that in *ex vivo* murine-derived enteroid models, hFcRn expression is necessary and sufficient for echovirus infection of the intestinal epithelium.

### Human FcRn is necessary but not sufficient for echovirus infection of the intestine *in vivo*

As described above, *ex vivo* enteroid models suggested that echovirus infection of murine-derived intestinal cells depended on expression of hFcRn and were not controlled by either type I or III IFNs. However, enteroids may not fully recapitulate the events associated with infection *in vivo*. To address this, we used six genotypes of mice, including the humanized FcRn models described above (hFcRn^Tg32^, hFcRn^Tg32^-IFNAR^-/-^, and hFcRn^Tg32^-IFNLR^-/-^) and animals expressing murine FcRn that were immunocompetent (C57/BL6, WT) or deficient in type I or III IFN signaling (IFNAR^-/-^ or IFNLR^-/-^, respectively) (**Figure 2A**). Neonatal (7-day old) mice were orally inoculated with 10^6^ PFU of E5 and monitored daily for 7 days for signs of illness (e.g., inactivity, discoloration, lack of nursing, lack of parental care, and death). We observed death in approximately 50% of hFcRn^Tg32^-IFNAR^-/-^ animals by 3 days post-inoculation (dpi) and almost 100% lethality by 7dpi (**Figure 2B**). In contrast, there were no clinical symptoms of illness in any other genotype and all animals survived until 7dpi (**Figure 2B**). There were no significant differences in mortality between male and female hFcRn^Tg32^-IFNAR^-/-^ mice (**Supplemental Figure 1A**).

**Figure 2.**
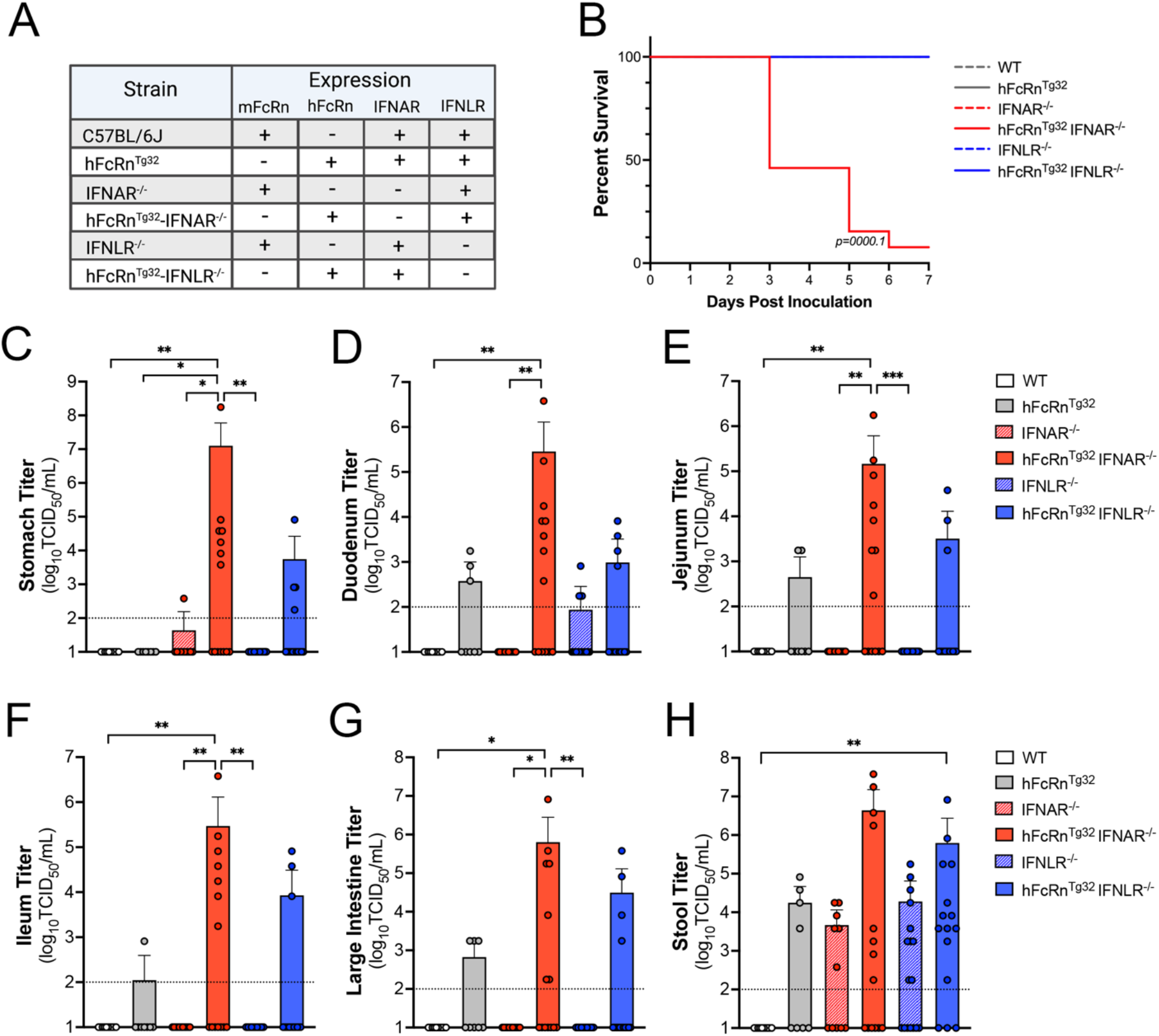
Expression of human FcRn is not sufficient for echovirus infection by the enteral route *in vivo*. **(A)** Table of the six genotypes used in this study. Shown is the expression of mouse or human FcRn, IFNAR, and IFNLR amongst these genotypes. **(B)** Survival of the indicated genotype of mice inoculated with 10^6^ E5 by oral gavage for 7 days post-inoculation. The log-rank test was used to analyze the statistical difference of the survival rate. **(C-H)**. At 3dpi, animals were sacrificed and viral titers in stomach **(C)**, duodenum **(D)**, jejunum **(E)**, ileum **(F)**, large intestine **(G)**, and stool **(H)** determined by TCID50 assays. In all, titers are shown as log_10_TCID50/mL with the limit of detection indicated by a dotted line. Data are shown as mean ± standard deviation with individual animals shown as each data point. Data are shown with significance determined with a Kruskal-Wallis test with a Dunn’s test for multiple comparisons (*p<0.05, **p<0.005, ***p<0.0005).

We next determined the extent of viral replication in the GI tract of infected animals at 3dpi by measuring viral titers in stomach, small intestine (duodenum, jejunum, and ileum), large intestine, and stool. Infected hFcRn^Tg32^-IFNAR^-/-^ animals contained high levels of virus in all tissues collected, with half or more of animals having high viral loads in stomach (7 of 14 mice), duodenum (8 of 14 mice), jejunum (8 of 14 mice), ileum (7 of 14 mice), and large intestine (7 of 14 mice) (**Figure 2C-H**). In contrast, there was no detectable virus in any tissues isolated from mice not expressing hFcRn, including WT (0 of 10 mice), IFNAR^-/-^ (0 of 11 mice), and IFNLR^-/-^ (0 of 17 mice) (**Figure 2C-H**). There were low levels of virus detected in select tissues from immunocompetent hFcRn^Tg32^ animals, which included duodenum (3 of 8 mice), jejunum (2 of 8 mice), ileum (1 of 8 mice), and large intestine (3 of 8 mice). No virus was recovered from the stomachs of hFcRn^Tg32^ mice (0 of 8 mice). Similarly, low to mid-level titers were observed in hFcRn^Tg32^-IFNLR^-/-^ mice, with virus recovered from stomach (3 of 15 mice), duodenum (4 of 15 mice), jejunum (3 of 15 mice), ileum (3 of 15 mice), and large intestine (4 of 15 mice). Stool collected from all genotypes except for WT contained high levels of virus in stool, which may reflect remaining inoculum. There were no significant differences in titers between male and female hFcRn^Tg32^-IFNAR^-/-^ mice, although male mice did have overall higher titers in various regions of the small intestine (**Supplemental Figure 1B**). These data show that hFcRn is necessary, but not sufficient, for echovirus infection of the intestine *in vivo* and that type I and III IFNs differentially control replication and pathogenesis.

### Type I IFNs are the primary drivers of dissemination outside of the GI tract

Given the high degree of mortality in orally inoculated hFcRn^Tg32^-IFNAR^-/-^ mice, we next assessed the levels of infection at key secondary sites of infection at 3dpi, including the liver, pancreas, and brain, which are all targeted by echoviruses in humans. hFcRn^Tg32^-IFNAR^-/-^ mice had higher levels of circulating virus (6 of 13 mice), which was not detected in any other genotype (**Figure 3A**). Consistent with this, we did not detect any virus in the livers, pancreases, or brains of WT, hFcRn^Tg32^, IFNAR^-/-^, IFNLR^-/-^, or hFcRn^Tg32^-IFNLR^-/-^ animals (**Figure 3B-D**). In contrast, hFcRn^Tg32^-IFNAR^-/-^ mice contained very high titers in liver (7 of 14 mice) and pancreas (7 of 14 mice) and lower titers in brain (5 of 14 mice) (**Figure 3B-D**). There were no significant differences in titers between male and female mice, although male mice did have overall higher titers in the liver (**Supplemental Figure 1B**). These data are consistent with our previous work data showing that hFcRn^Tg32^-IFNAR^-/-^ pups or adult mice inoculated by the IP route have high levels of echovirus infections in the liver and pancreas^21^.

**Figure 3.**
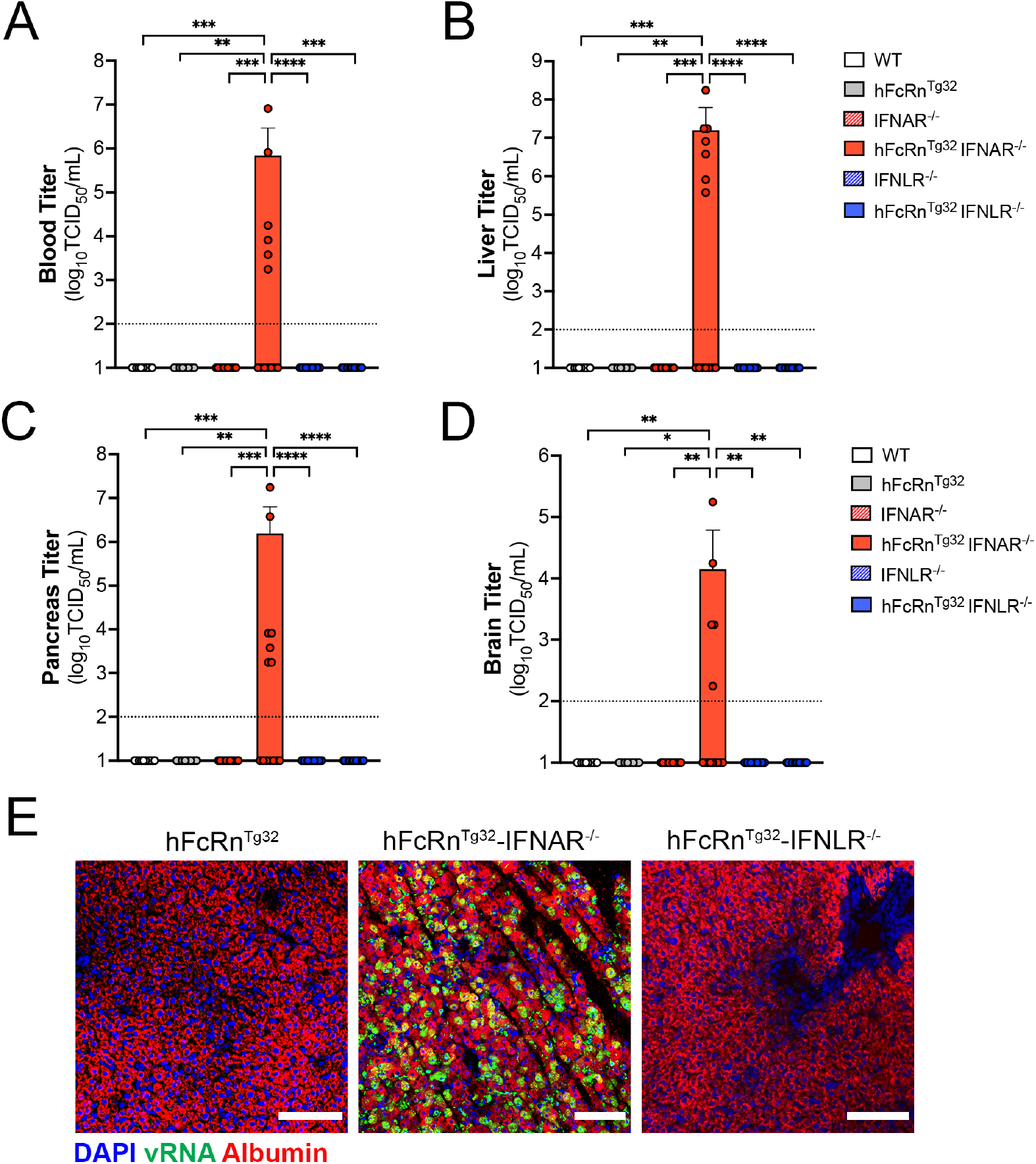
Type I IFNs control echovirus dissemination from the GI tract. 7-day old pups were orally inoculated with 10^6^ PFU of E5 and at 3dpi, sacrificed for viral titration and histology. **(A-D)**, Viral titers in the blood **(A)**, liver **(B)**, pancreas **(C)**, and brain **(D)** are shown. In all, titers are shown as log_10_TCID50/mL with the limit of detection indicated by a dotted line. Data are shown as mean ± standard deviation with individual animals shown as each data point. **(E)** Hybridization chain reaction RNA-FISH (HCR) from liver section of hFcRn^Tg32^, hFcRn^Tg32^-IFNAR^-/-^, or hFcRn^Tg32^-IFNLR^-/-^ neonatal mice at 3dpi using probes against the E5 genome (green) and albumin (red). DAPI-stained nuclei are shown in blue. Scale bars shown at bottom right (100χm). In A-D, data are shown with significance determined with a Kruskal-Wallis test with a Dunn’s test for multiple comparisons (*p<0.05, **p<0.005, ***p<0.0005, ****p<0.0001).

Next, we performed Luminex multiplex assays to determine the levels of twenty-five circulating cytokines in the blood of E5-infected animals. Consistent with their high levels of dissemination, we found that hFcRn^Tg32^-IFNAR^-/-^ mice induced pronounced antiviral and pro-inflammatory signaling in response to E5 infection, which included high levels of circulating type I IFNs (IFN-α and IFN-β), G-CSF, and IL-6 (**Supplemental Figure 2A-D**). No other genotypes contained any significant increases in circulating cytokines (**Supplemental Figure 2A-D**). These data are similar to our previous work where IP inoculated hFcRn^Tg32^-IFNAR^-/-^ animals had high levels of circulating type I IFNs^21^.

Because we observed significant dissemination of E5 to the livers of orally inoculated hFcRn^Tg32^-IFNAR^-/-^ mice, we next determined if the cellular tropism of echoviruses is the same between the IP and oral routes of inoculation. To do this, we performed hybridization chain reaction (HCR), which allows for multiplexed fluorescent quantitative RNA detection with enhanced sensitivity over conventional hybridization approaches^24,25^. Our previous work using this method showed that echoviruses exclusively target hepatocytes following IP inoculation^21^. We designed probes specific for the E5 genome and used probes to the hepatocyte marker albumin and performed HCR on liver sections from hFcRn^Tg32^, hFcRn^Tg32^-IFNAR^-/-^, and hFcRn^Tg32^-IFNLR^-/^ mice orally inoculated with E5 at 3dpi. E5 vRNA positive cells exclusively colocalized with albumin, identifying hepatocytes as the main cellular target of infection in the liver following dissemination from the GI tract (**Figure 3E**). Collectively, these data show that type I IFNs are the primary drivers of echovirus dissemination from the GI tract to secondary sites including the liver and pancreas.

### Type III IFNs limit persistent echovirus infection in the GI epithelium

Because we observed low levels of E5 replication in GI-derived tissues at 3 dpi, we next compared viral titers from tissues isolated at 7dpi to determine if there were differences in persistence compared to hFcRn^Tg32^-IFNAR^-/-^ mice. As hFcRn^Tg32^-IFNAR^-/-^ mice died from disease before 7dpi, they were excluded from these studies. At 7dpi, hFcRn^Tg32^-IFNLR^-/-^ mice were the only genotype with consistently detectable virus in tissues associated with the GI tract. Whereas select animals had detectable virus in the stomach (1 of 7 hFcRn^Tg32^ mice and 2 of 14 hFcRn^Tg32^-IFNLR^-/-^ mice) and large intestine (3 of 14 hFcRn^Tg32^-IFNLR^-/-^ mice), hFcRn^Tg32^-IFNLR^-/-^ animals had higher levels of virus in all regions of the small intestine including the duodenum (6 of 14 mice), jejunum (6 of 14 mice), and ileum (4 of 14 mice) (**Figure 4A-G**). Consistent with more persistent infection in the GI tract of hFcRn^Tg32^-IFNLR^-/-^ mice, these mice also contained higher levels of virus in stool (7 of 14 animals with detectable virus) compared to all other genotypes (**Figure 4F**). Male animals did contain higher viral titers than did female mice, although these differences were not significant (**Supplemental Figure 1C**). However, even at 7dpi, we were unable to detect any virus in the blood, liver, pancreas, or brain of any genotype, including hFcRn^Tg32^-IFNLR^-/-^ (**Supplemental Figure 3A, B, C, & D)**. These data show that type III IFNs do not control dissemination but limit persistent infection of the intestinal epithelium.

**Figure 4.**
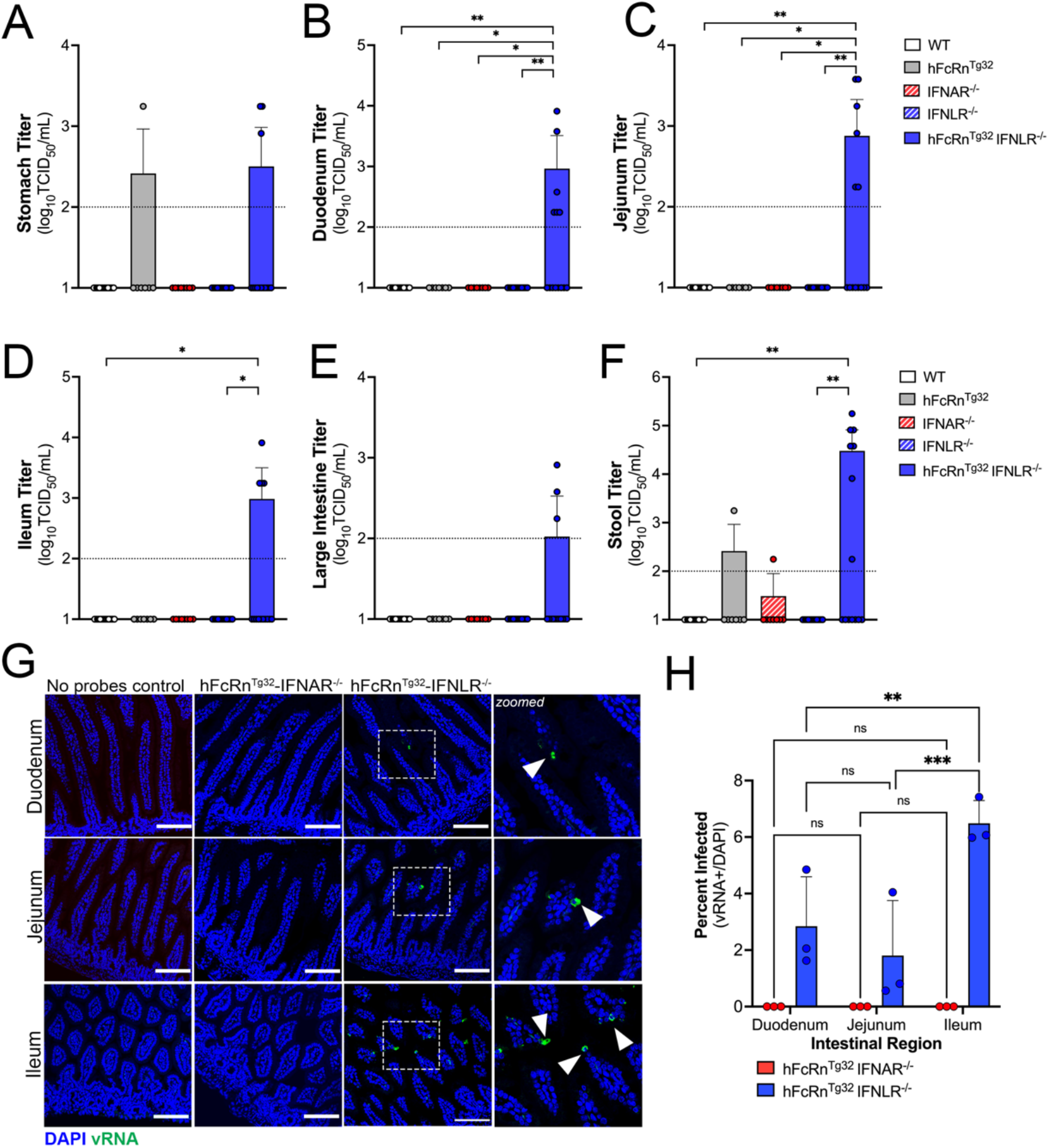
Type III IFNs restrict persistent echovirus infection in the GI epithelium. 7-day old neonatal were orally inoculated with 10^6^ PFU of E5 and at 7dpi, animals were sacrificed for viral titration and tissue collection. **(A-F)**, Viral titers are shown in stomach **(A)**, duodenum **(B)**, jejunum **(C)**, ileum **(D)**, large intestine **(E)**, and stool **(F)**. In all, titers are shown as log_10_TCID50/mL with the limit of detection indicated by a dotted line. Data are shown as mean ± standard deviation with individual animals shown as each data point. Significance was determined using a Kruskal-Wallis test with a Dunn’s test for multiple comparisons (*p<0.05, **p<0.005). **(G)** At 3dpi, animals were sacrificed and the entire GI tract was removed and swiss rolled following by histologic sectioned. HCR of hFcRn^Tg32^-IFNAR^-/-^ or hFcRn^Tg32^-IFNLR^-/-^ pups at the 3dpi using probes against the E5 genome (green) and DAPI (blue). Scale bars shown at bottom right (100χm). Zoom of specific regions in hFcRn^Tg32^-IFNLR^-/-^ images are shown to the right. **(H)** Quantification of three independent tile scans using confocal microscopy of each region of the small intestines based on the number of villi that were positive for vRNA using the cell count function in FIJI. Data are shown as percent of vRNA positive villi over total villi per tile scan. Three independent tile scans were quantified (for an average of 144 villi in the duodenum, 224 villi in the jejunum, and 164 villi in the ileum). Significance was determined by a Two-way Anova with Šídák’s multiple comparisons tests (*p<0.05,**p<0.005, ***p<0.0005, ****p<0.0001).

Visualization of intestinal replication of enteroviruses *in vivo* has been hindered by the lack of sensitive assays to monitor infection with low signal-to-noise. To overcome this limitation, we utilized HCR, a component of which includes signal amplification given the self-assembly of secondary detection hairpins into amplification polymers. We inoculated 7-day old hFcRn^Tg32^-IFNAR^-/-^ and hFcRn^Tg32^-IFNLR^-/-^ animals with 10^6^ PFU of E5 by the oral route, sacrificed them at 3dpi, and then performed Swiss rolling of full intestinal tissue, which was sectioned and processed for HCR. In contrast to viral titer data, which showed high levels of virus in the intestines of hFcRn^Tg32^-IFNAR^-/-^ animals, we did not detect any vRNA in any intestinal section of these animals **(Figure 4G)**. However, we observed clear areas of vRNA-containing cells in various regions of the small intestines of hFcRn^Tg32^-IFNLR^-/-^ animals **(Figure 4G)**. While vRNA was detected in both the duodenum and jejunum, there were more vRNA-containing cells in the ileum of the hFcRn^Tg32^-IFNLR^-/-^ animals **(Figure 4G, 4H)**, suggesting that there may be regional differences in echovirus persistence in the epithelium. Although we observed areas of viral replication within the epithelium, we do not see any damage to the epithelium. A blinded pathologist reviewed H&Es from uninfected, hFcRn^Tg32^-IFNAR^-/-^ and hFcRn^Tg32^-IFNLR^-/-^ animals and observed no significant changes or damage to the intestine following 3dpi E5 infection (**Supplemental Figure 4A**). Additionally, we observed no change to the cellular composition of the epithelium as suggested by Periodic Acid Schiff (PAS) staining for goblet cells (**Supplemental Figure 4B**)

### Enterocytes are the main cellular targets of echoviruses *in vivo*

We showed previously that echoviruses preferentially infect enterocytes and enteroendocrine cells in human stem cell-derived enteroids^9^. However, whether there is a cell type specificity of infection for echoviruses, or other enteroviruses, *in vivo* is unknown. To define the cellular tropism of echoviruses *in vivo*, we designed HCR probes targeting an enterocyte marker (alkaline phosphate intestinal, Alpi), goblet cell marker (mucin-2, Muc2), and enteroendocrine cell marker (chromogranin A, Chga). We confirmed the specificity of these probes in murine-derived intestinal tissue and found that they accurately labeled distinct cell populations in the epithelium (**Figure 5A**). Using probes directed against E5 and Alpi, Muc2, or Chga, we performed HCR in Swiss rolled intestinal tissue sections isolated from hFcRn^Tg32^-IFNLR^-/-^ animals orally infected with 10^6^ E5 at 3dpi. We found that echovirus vRNA exclusively localized to Alpi-positive cells (**Figure 5B**) and was not observed in any Muc2-positive goblet cells (**Figure 5C**) or Chga-positive enteroendocrine cells (**Figure 5D**), as assessed by image analysis and quantification of confocal microscope-generated tile scans of ∼4mm^2^ of intestinal tissue (**Figure 5E and Supplemental Figure 5**). These data show that enterocytes are the main targets of echoviruses following oral inoculation of hFcRn^Tg32^-IFNLR^-/-^ mice.

**Figure 5.**
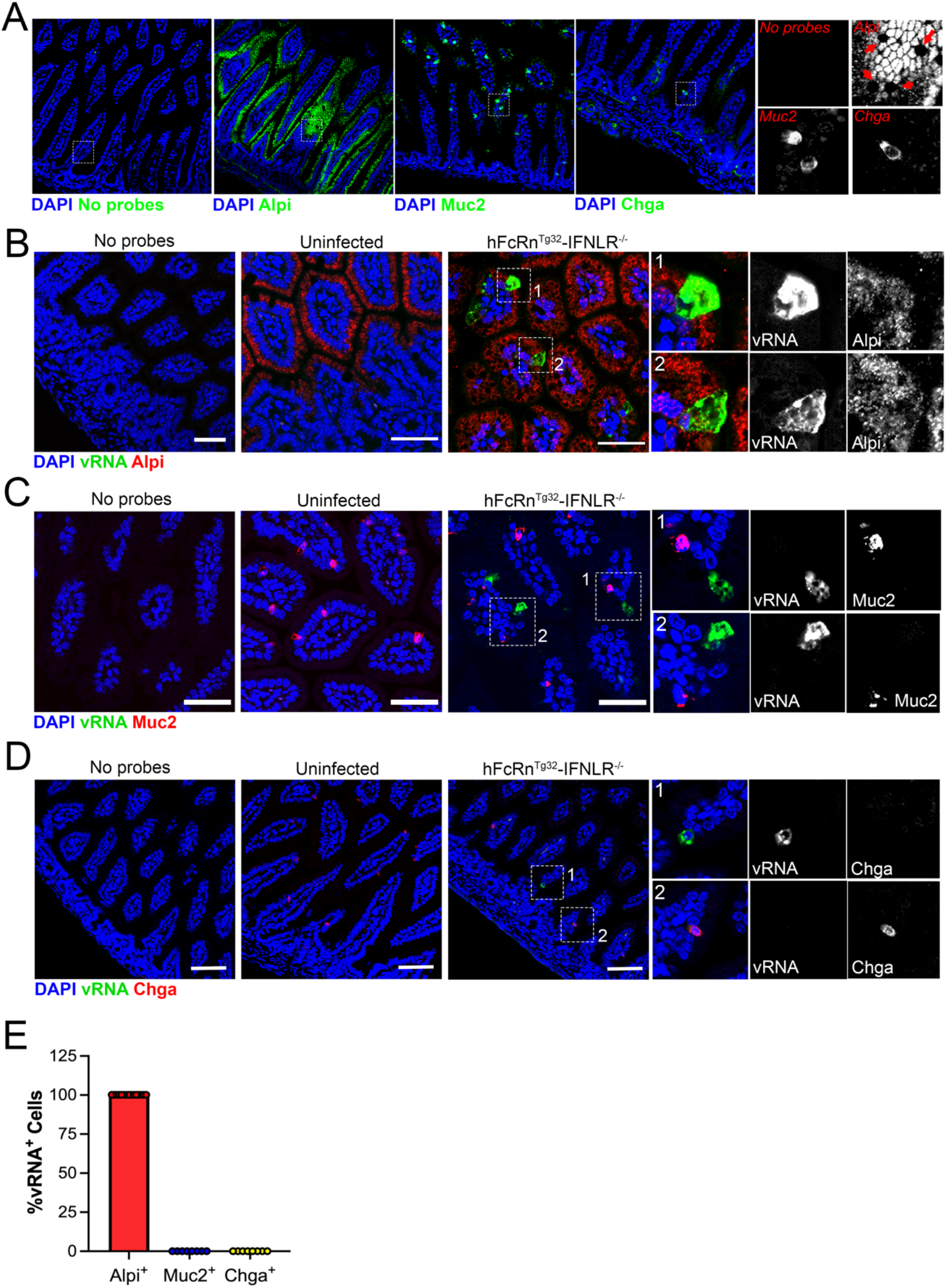
*In vivo* replication of echoviruses is specific for enterocytes. **(A)** Hybridization chain reaction RNA-FISH (HCR) of uninfected small intestine sections using specific probes against Alpi, Muc2, or Chga (in green), as indicated at bottom. DAPI-stained nuclei are shown in blue. No probe containing control is shown at left. In all, white box is shown zoomed (∼6x) at right using the probes indicated in red. Red arrows in Alpi section denote goblet cells based on morphology that were not positive for Alpi, as expected. **(B-D)** 7-day old hFcRn^Tg32^-IFNLR^-/-^ neonatal mice were orally inoculated with 10^6^ PFU of E5 and at 3dpi, animals were sacrificed and the entire small intestine removed and Swiss rolled for subsequent histologic sectioning. Shown are representative images of ileum tissue using probes to E5 (green in all) and either Alpi (B), Muc2 (C), or Chga (D) (red in all). DAPI-stained nuclei are shown in blue. In all, white boxes denote zoomed areas shown at right, which include black and white images as indicated. Scale bars shown at bottom right (50χm). **(E)** Quantification of confocal images was performed using Fiji and was quantified as the total percentage of vRNA positive cells that colocalized with Alpi (in red), Muc2 (in blue), or Chga (in green). Note that there was no colocalization between vRNA and either Muc2 or Chga.

## Discussion

The events associated with enterovirus infections of the GI tract *in vivo* are largely unknown. Here, we defined the role of hFcRn and type-specific IFN signaling in mediating echovirus infections of the intestinal epithelium and dissemination to secondary tissue sites. We show that hFcRn is necessary and sufficient for echovirus infection of the intestinal epithelium in enteroids derived from humanized FcRn mice. However, *in vivo*, expression of hFcRn alone is not sufficient for echovirus infection by the enteral route. Using humanized FcRn mouse models deficient in either type I or III IFN signaling, we defined the differential roles of these IFNs in echovirus replication in and dissemination from the GI tract. These studies showed that type I IFNs limit dissemination of echoviruses from the GI tract and ablation of this signaling robustly increases viral replication at secondary sites, such as the liver. In contrast, type III IFNs suppress replication in the intestinal epithelium and deletion of the receptor for these IFNs prolongs intestinal echovirus replication and increases viral persistence. We further show that echoviruses preferentially infect enterocytes *in vivo*, which is enhanced in the absence of type III IFN signaling. Collectively, our work presented here provides key insights into the roles of FcRn and IFN signaling in echovirus pathogenesis in the GI tract.

Little is known regarding the mechanisms used by echoviruses to enter the intestinal epithelium. Our data support a model whereby hFcRn is necessary and sufficient for intestinal replication *in vitro*. While some echoviruses utilize decay accelerating factor (DAF/CD55) as an attachment factor *in vitro*^26^, E5 does not bind DAF^19^. Moreover, DAF-binding echoviruses do not bind the murine homolog of DAF^26^. While a previous study predicted that echovirus binding to DAF might trigger viral internalization and particle delivery to endosomes, at which time FcRn-mediated uncoating would occur^20^, the data presented here do not support such a model and suggest that DAF plays no role in echovirus infections of the intestinal epithelium *in vitro* or *in vivo*. Instead, our data suggest that FcRn is necessary and sufficient for echovirus infection of the intestinal epithelium and occurs independent of DAF binding. This is consistent with *in vivo* data from humanized mouse models of DAF, which show that expression of DAF does not impact intestinal replication of DAF-binding variants of CVB^14^.

FcRn is unique in its ability to mediate the transcytosis of IgG and albumin across the intestinal epithelium. Interestingly, this transport functions in a bidirectional manner in cultured intestinal cell lines, suggesting that FcRn can sample contents from the apical or basolateral domains and mediate the transcytosis of cargo to the opposing domain^27^. This function of FcRn could have important implications during echovirus infections—FcRn could *(i)* mediate the internalization of viral particles into intracellular compartments that facilitate uncoating and subsequent replication and/or *(ii)* could mediate the direct transcytosis of viral particles across the intestinal epithelium from the lumen into underlying tissue. Given that FcRn mediates bidirectional transport across the epithelium, this raises the possibility that echoviruses could be transported from either the apical or basolateral domains to cross the intestinal barrier. We were unable to visualize active replication in the intestinal epithelium of hFcRn^Tg32^-IFNAR^-/-^ animals, despite robust viral dissemination to secondary sites of infection. In contrast, we detected vRNA in ∼6% of enterocytes in ileum tissue of hFcRn^Tg32^-IFNLR^-/-^ animals, in which there was no dissemination observed. These data suggest that in addition to facilitating viral entry and replication into enterocytes, it is possible that in some cases, FcRn might facilitate the transcytosis of echovirus particles across the epithelium and that ablation of type I IFN signaling promotes dissemination of these particles to secondary sites of infection.

Type III IFNs are important in antiviral defenses of many barrier tissues, including the GI tract^28,29^. For example, IFN-λs control rotavirus infection in the intestinal epithelium in adult and neonatal mice^17^. This study showed that whereas mice lacking IFNLR were more susceptible to rotavirus replication and viral-induced cytotoxicity, IFNAR^-/-^ mice were comparable to immunocompetent WT mice. These data are distinct from our work presented here, which shows that type I IFNs are key host mediators that prevent echovirus dissemination following oral infection. Type III IFNs have also been implicated in restricting murine norovirus replication in the GI tract *in vivo*^16^. Similar to our findings with echoviruses, IFN-λs restrict persistent norovirus infection whereas type I IFNs restrict dissemination^16^. Previous studies with PV and EV71 suggest that type I IFNs control viral replication of these enteroviruses by the enteral route *in vivo*^12,13^ whereas CVB infection is unchanged in animals deficient in IFNAR^14^. While the mechanistic basis for these differences is unknown, it is possible that the cell-type specific nature of enterovirus replication in the intestine may influence their dependence on IFN signaling. For example, in human enteroids, EV71 preferentially infects goblet cells whereas echoviruses are enriched in enterocytes^9,10^. In cell lines, previous work has suggested that PV transcytoses across M cells, suggesting it does not replicate in the epithelium^30^. Future studies on the cell-type specific nature of IFN signaling in distinct lineages of intestinal cells and the impact of these differences on enterovirus replication will be essential to determine if the distinct cellular tropism of enteroviruses in the GI tract influences IFN-mediated signaling.

Our findings presented here define fundamental aspects of echovirus biology that enhance our understanding of how infection, tissue targeting, and disease occurs *in vivo*. We show that FcRn is necessary but not sufficient for echovirus infections of the GI tract in vivo and that type I and III IFNs differentially control echovirus persistence and dissemination. Collectively, these studies provide new insights into echovirus biology and the development of in vivo models that recapitulate distinct aspects of echovirus disease, which could potentially accelerate the development of therapies.

## Materials and Methods

### Cell lines and viruses

HeLa cells (clone 7B) were provided by Jeffrey Bergelson, Children’s Hospital of Philadelphia, Philadelphia, PA, and cultured in MEM supplemented with 5% FBS, non-essential amino acids, and penicillin/streptomycin. Experiments were performed with echovirus 5 (Noyce strain, E5), which was obtained from the ATCC. Virus was propagated in HeLa cells and purified by ultracentrifugation over a 30% sucrose cushion, as described previously^31^. Enteroid experiments were performed with light-sensitive neutral red (NR) incorporated viral particles. E5 was propagated in the presence of NR (10μg/mL) in semi-dark conditions and was subsequently purified in semi-dark conditions by ultracentrifugation over a sucrose cushion^32^. All viruses were sequenced for viral stock purity following propagation. Purity of all viral stocks was confirmed by Sanger sequencing of VP1 using enterovirus-specific primers^33^. Briefly, RNA extraction was performed on 10μl of purified virus stock, according to manufacturer’s instructions (Qiagen Cat. 529904). RNA was reverse transcribed using SuperScript III reverse transcription kit, (Invitrogen cat. 18080093) according to manufacturer’s instructions, with a pan enterovirus primer (vir21; ATAAGAATGCGGCCGCTTTTTTTTTTTTTTTTTTTTTTTTT), followed by an RNaseH treatment for 20 minutes. PCR was performed with 5μl of the cDNA reaction using BioRad iTaq DNA polymerase (BioRad cat. 1708870). Virus specific primers were as follows: E5 forward 5’-TATCGCCAATTACAACGCGAA-3’; E5 reverse 5’-TTGGTTTGAAGTAAACCCTTA-3’.

### Animals

All animal experiments were approved by the Duke University Animal Care and Use Committees, and all methods were performed in accordance with the relevant guidelines and regulations. C57BL/6J (WT, cat. no. 000664), B6.Cg-*Fcgr*^*t*tm1Dcr^Tg(FCGRT)32Dcr/DcrJ (hFcRn^Tg32^, cat. no. 014565), and B6.(Cg)-Ifnar1^tm1.2Ees^/J (IFNAR^-/-^, cat. no. 028288) mice were purchased from The Jackson Laboratory. hFcRn^Tg32^-IFNAR^-/-^ mice were generated as described previously^21^. B6.Ifnlr^-/-^/J (IFNLR^-/-^) mice were provided by Dr. Megan Baldridge (Washington University School of Medicine). hFcRn^Tg32^-IFNLR^-/-^ mice were generated by crossing B6.Cg-*Fcgr*^*t*tm1Dcr^Tg(FCGRT)32Dcr/DcrJ (hFcRn^Tg32^, cat. no. 014565) mice with B6.Ifnlr^-/-^/J mice. Breeders were established that were deficient in mouse FcRn and IFNLR and were homozygous for the hFcRn transgene. All animals used in this study were genotyped by Transnetyx.

### Enteroid isolation and passaging

Murine intestinal crypts were isolated using a protocol adapted from Stem Cell Technologies. Briefly, intestines were isolated from five 10-day old pups and connective tissue removed. Intestines were cut longitudinally and washed extensively in PBS. Intestines were cut into 5mm segments and washed again using a 10mL serological pipette until PBS was clear. Washed intestinal pieces were incubated in gentle cell dissociation reagent (Stem Cell Technologies Cat. 07174) for 15 minutes at room temperature. Crypts were released using 0.1% BSA and vigorous pipetting with a 10mL serological pipette. Crypts were filtered using a 70μm cell strainer and the resulting flow through centrifuged at 290xg for 5 minutes. Pellets were resuspended in Matrigel (Corning Cat. 356231) and 40μL of crypt-containing Matrigel ‘domes’ plated into each well of a 24 well plates (Corning 3526), placed in a 37°C incubator to pre-polymerize for ∼3 min, turned upside-down to ensure equal distribution of the isolated cells in domes for another 10 min, then carefully overlaid with 500 μL IntestiCult Organoid Growth Medium (Stem Cell Technologies Cat. 06005) supplemented with 1% penicillin/streptomycin, 50μg/mL gentamycin, and 0.2% amphotericin b, containing Y-27632 (Rock inhibitor, Sigma). Media was changed every 48hrs and Y-27632 was removed after the first media change.

Confluent enteroids were passaged by manual disruption of Matrigel domes with a P1000 pipette tip in PBS and centrifuged at 400xg for 5 minutes. The enteroid-containing pellet were resuspended in TrypLE (Invitrogen Cat. 12605010) and incubated in a water bath at 37°C for 8 minutes. Enzyme activity was quenched with DMEM containing 10% FBS and centrifuged at 400xg for 5 minutes. Pellets were resuspended in Matrigel and 40μL Matrigel domes in each well of a 24 well plate. Domes were allowed to solidify at 37°C for 10 minutes and then covered with IntestiCult Organoid Growth Medium, as described above. For infections, crypts were plated in 24-well plates pre-coated with 15uL of Matrigel using a P1000 tip and allowed to solidify for 30 minutes.

### Enteroid Infections

Enteroids plated on Matrigel coating as described above were allowed to differentiate for 5 days with media replaced every 48hrs. For infections, wells were infected with 10^6^ PFU of NR-incorporated virus, generated as described above. Virus was pre-adsorbed for 1 hour at 16°C, enteroids washed three times with PBS, and media replaced. Infections were initiated by shifting enteroids to 37°C. At 6hpi, enteroids were exposed to light on a light box for 20 minutes to render intact viral particles non-infectious and infections performed for the times indicated. Plaque assays were performed in HeLa cells overlayed with 1:1 mixture of 1% agarose and 2x MEM (4% FBS, 2% pen/strep, 2% NEAA). Plaques were enumerated 40hpi by crystal violet staining.

### RNA extraction and RNAsequencing

Total RNA was prepared using the Sigma GenElute total mammalian RNA miniprep kit with optional DNase step, according to the protocol of the manufacturer. RNA quality was assessed by Nanodrop and an Agilent RNA Screen Tape System, and 1ug was used for library preparation using RNA with Poly A selection kit (Illumina), as per the manufacturer’s instructions. Sequencing was performed on an Illumina HiSeq. RNA-seq FASTQ data were processed and mapped to the mouse reference genome (GRCm38) using CLC Genomics Workbench 20 (Qiagen). Differential gene expression was performed using the DESeq2 package in R^34^. Heatmaps and volcano plots were made in GraphPad Prism 9. Raw sequencing files have been deposited in Sequence Read Archives.

### Suckling pup infections

7-day-old mice were inoculated by the oral route with 10^6^ PFU of E5. Oral gavage inoculation was performed using a 1mL disposable syringe and a 24-gauge round tipped needle in 50μL of 1X PBS. Mice were euthanized at either 3- or 7-days post inoculation and organs harvested into 0.5mL of DMEM and stored at -80°C. Tissue samples for viral titration were thawed and homogenized with a TissueLyser LT (Qiagen) for 5 minutes, followed by brief centrifugation for 5 minutes at 8000xg. Viral titers in organ homogenates were determined by TCID50 in HeLa cells and enumerated following crystal violet staining.

### HCR and Imaging

HCR was performed following the Molecular Instruments HCR v3.0 protocol for FFPE human tissue sections^24,25^. Briefly, tissue sections were deparaffinized with xylene and rehydrated with decreasing concentrations of ethanol (100%, 95%, 80%). Antigen unmasking was performed with slides submerged in 10 mM citrate buffer (pH 6.0) and heated in a steamer for 20 minutes at ∼90^°^C. Slides were cooled to room temperature. Sections were treated with 10 µg/mL Proteinase K for 10 min at 37^°^C and washed with RNase free water. Samples were incubated for 10 minutes at 37^°^C in hybridization buffer. Sections were incubated overnight in a humidified chamber at 37^°^C with 3 pmol of initiator probes in hybridization buffer. We designed probes for E5 (**Supplemental Table 1**), Muc2 (**Supplemental Table 2**), and Chga (**Supplemental Table 3**) in house. Custom probes for Alpi were designed by Molecular Instruments (Lot PRI910). The next day, slides were washed in probe wash buffer and 5x SSCT for 4x 15 min, according to the manufacturer’s instructions. Samples were incubated in a humidified chamber at 37^°^C for 30 minutes in amplification buffer. Fluorescent hair pins were heated to 95^°^C for 90 seconds and snap cooled at room temperature for 30 min. Hairpins and amplification buffer were added to the sample and incubated overnight at room temperature. Hairpins were washed off with 5x SSCT for 5 minutes, 15 minutes, 15 minutes, and 5 minutes followed by a wash with PBS containing DAPI. Slides were mounted in vectashield with DAPI. Slides were imaged on a Zeiss 880 with Airyscan inverted confocal microscope. Tile scans were performed at a 20x magnification using a 6 by 6 square area resulting in 36 total images. Each intestinal segment was tile scanned using three different areas for quantification. Image analysis was performed using FIJI.

### Periodic Acid Schiff (PAS) staining

PAS staining was performed according to manufactures instructions (Abcam, ab150680). Slides were mounted with Cytoseal 60 (Thermo Scientific, 83104). Images were captured on an IX83 inverted microscope (Olympus) using a UC90 color CCD camera (Olympus).

### Statistics

All statistical analysis was performed using GraphPad Prism version 8. Data are presented as mean ± SD. A one-way ANOVA was used to determine statistical significance, as described in the figure legends. Parametric tests were applied when data were distributed normally based on D’Agostino–Pearson analyses; otherwise, nonparametric tests were applied. P values of <0.05 were considered statistically significant, with specific P values noted in the figure legends.

## Acknowledgements

We thank Cristian Ovies for technical assistance (Duke University), Megan Baldridge (Washington University) for providing IFNLR^-/-^ mice, and Sujan Shresta (La Jolla Institute for Immunology) for providing hFcRn^Tg32^-IFNAR^-/-^ mice. This project was supported by NIH R01-AI150151 (C.B.C), NIH T32-AI060525 (A.I.W), NIH F31-AI149866 (A.I.W). The funders had no role in study design, data collection and analysis, decision to publish, or preparation of the manuscript.

---

**Supplemental Table 1.**
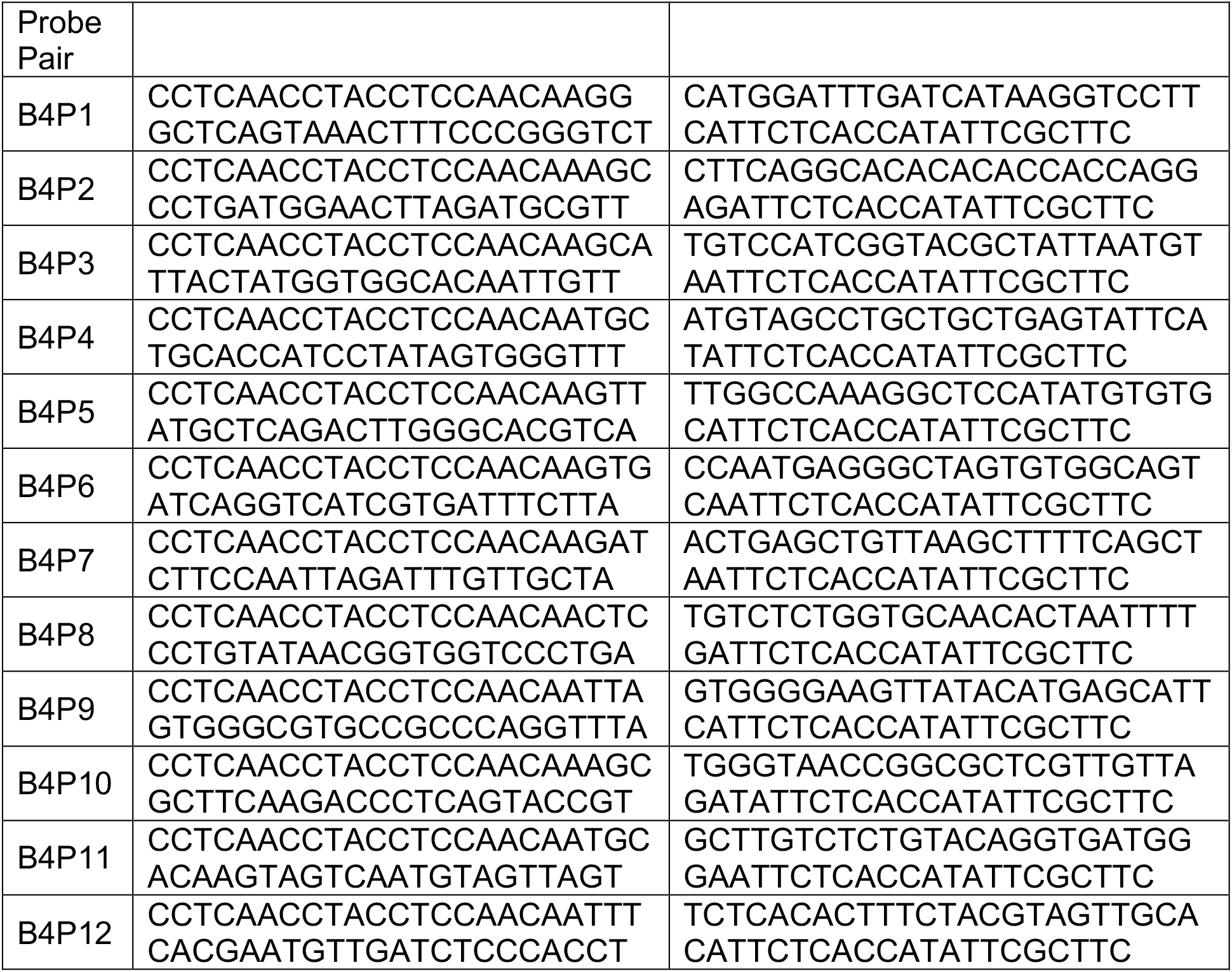
Echovirus 5 HCR Probes.

**Supplemental Table 2.**
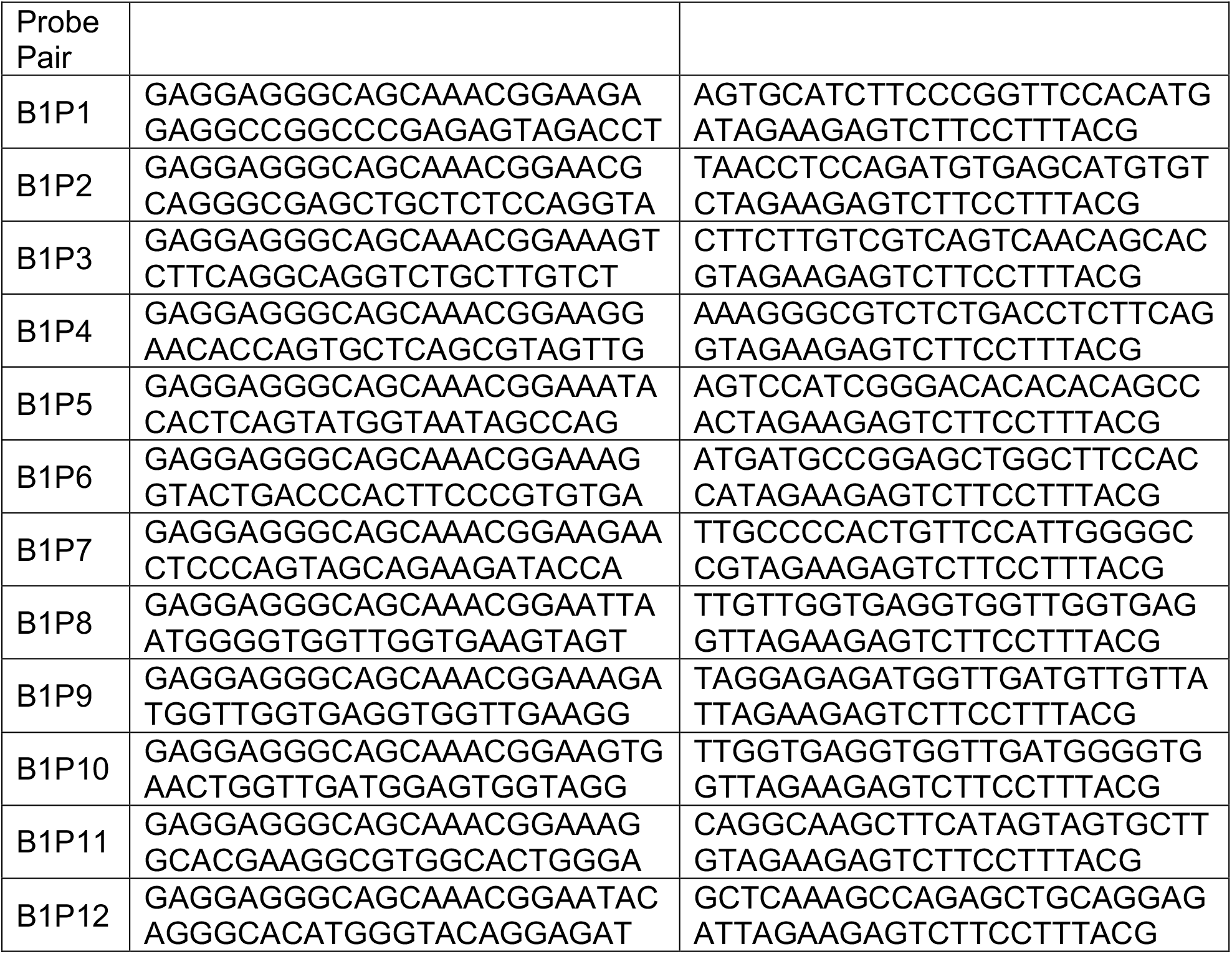
Muc2 HCR Probes.

**Supplemental Table 3.**
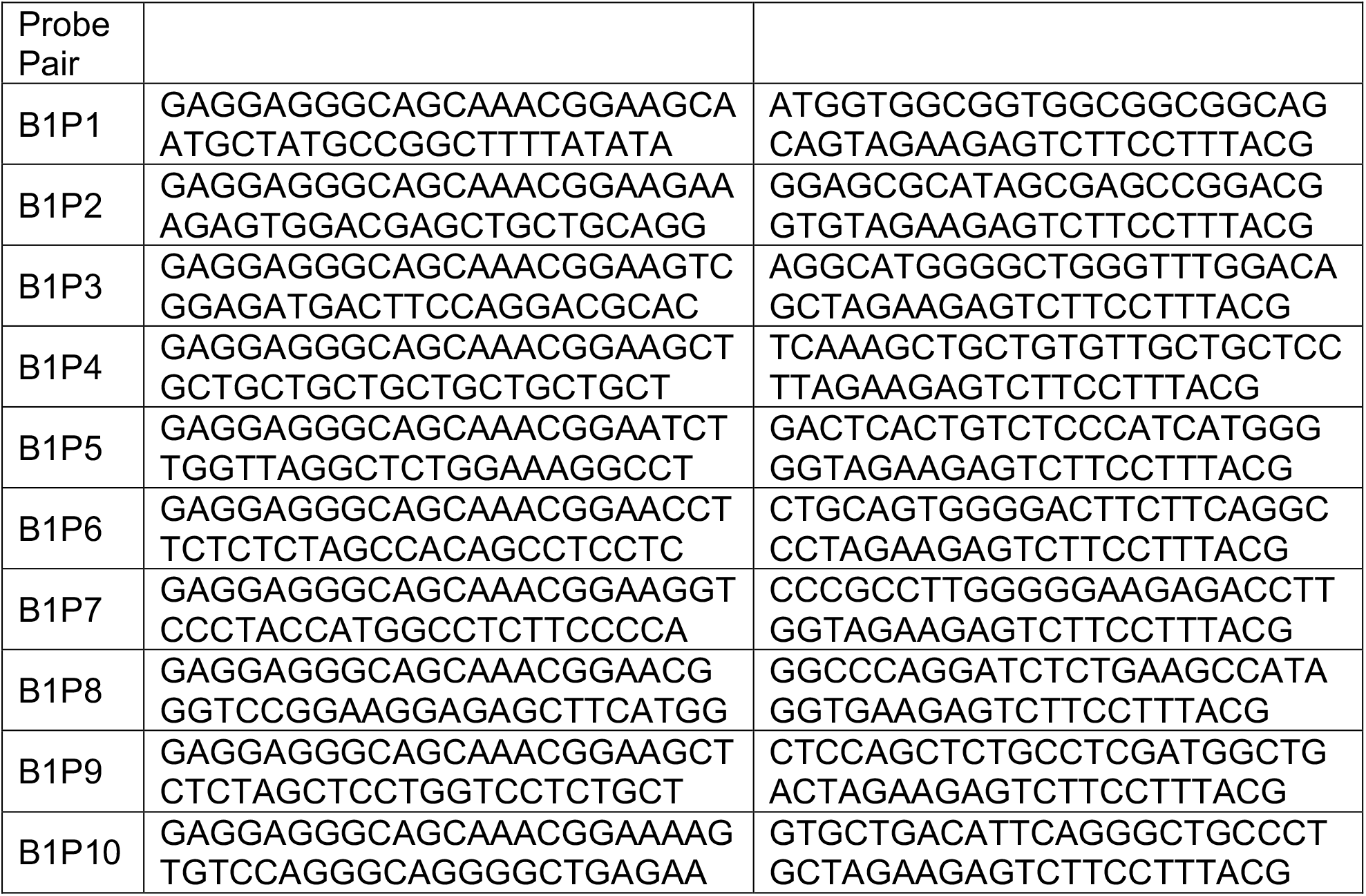
Chga HCR Probes.

**Supplemental Figure 1.**
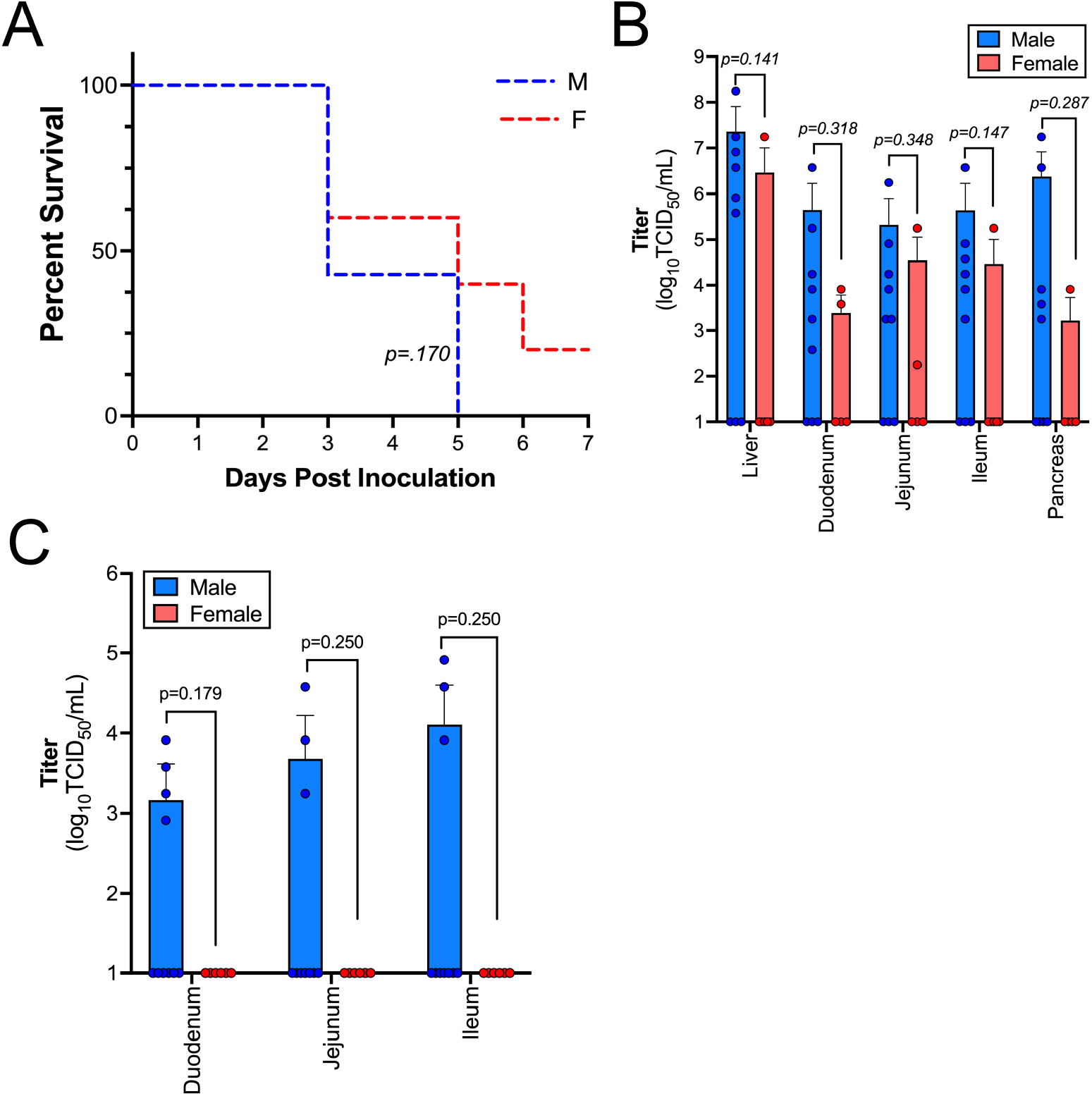
7-day old pups were orally inoculated with 10^6^ PFU of E5. **(A)** Survival curve of hFcRn^Tg32^-IFNAR^-/-^ animals broken down by sex. A log-rank test was used to analyze the statistical difference of the survival rate. **(B)** Viral titers of hFcRn^Tg32^-IFNAR^-/-^ animals at 3dpi broken down by sex. **(C)** Viral titers of hFcRn^Tg32^-IFNLR^-/-^ animals at 7dpi broken down by sex. Significance was determined by Mann-Whitney U test (p values shown). Each symbol represents an individual animal.

**Supplemental Figure 2.**
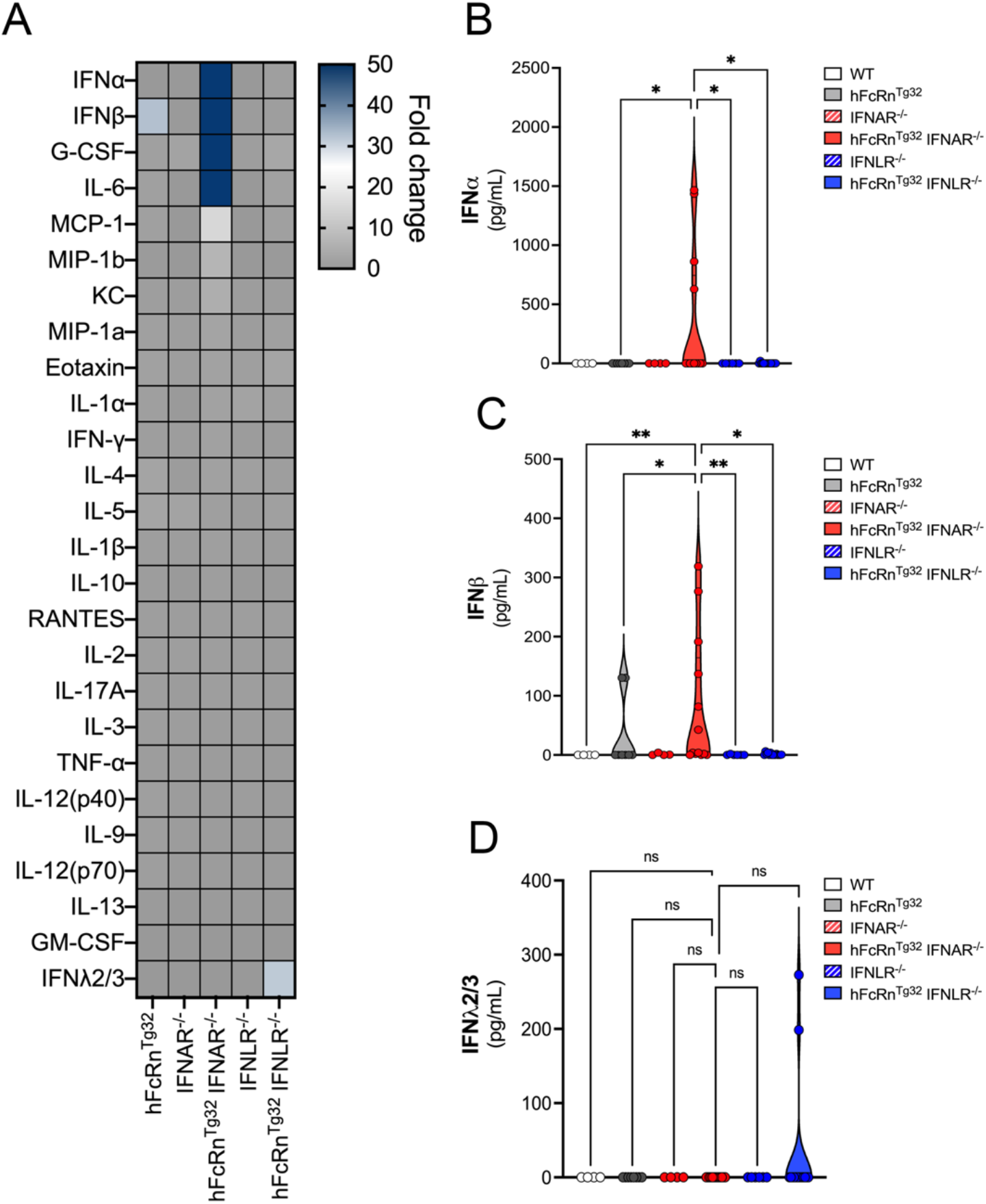
Neonatal mice were inoculated by the oral route with 10^6^ PFU of E5 and sacrificed 3 dpi. Luminex-based multianalyte profiling of 26 cytokines was then performed from whole blood. **(A)** Heatmap demonstrating the induction (shown as fold-change from uninfected control) in E5-infected mice of the indicated genotype. Blue denotes significantly increased cytokines in comparison to untreated. Grey or white denote little to no changes (scale at top right). The IFNs are shown to the right as pg/mL IFNα **(B)**, IFNβ **(C)**, and IFNλ2/3 **(D)**. Data are shown as mean ± standard deviation and individual animals (points). Data are shown with significance determined with a Kruskal-Wallis test with a Dunn’s test for multiple comparisons (*p<0.05, **p<0.005, ns-nt significant). Each symbol represents an individual animal.

**Supplemental Figure 3.**
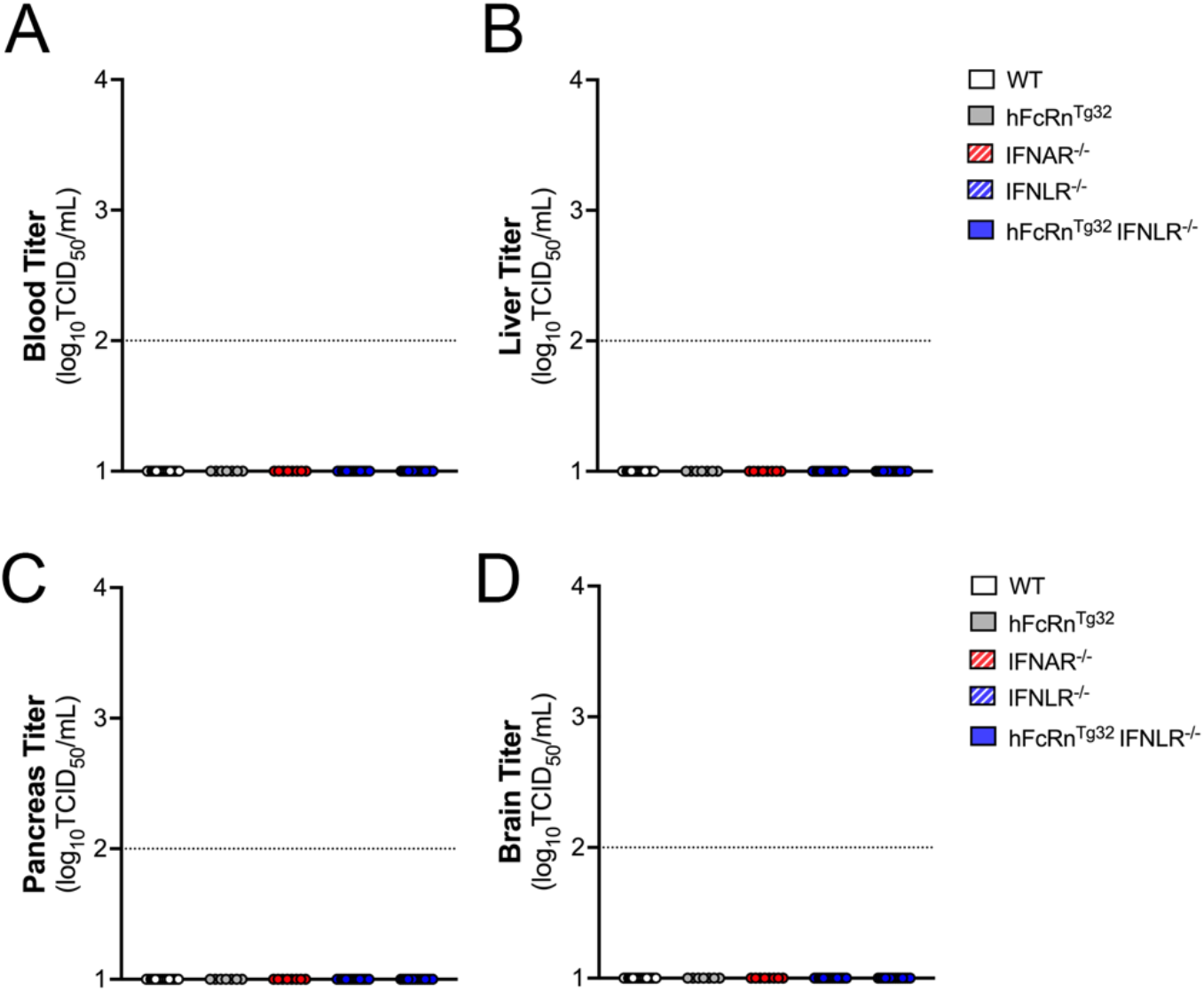
7-day old pups were orally inoculated with 10^6^ PFU of E5. At 7dpi, animals were sacrificed to measure viral replication in tissues. Viral titers are shown as log_10_TCID50/mL in the blood **(A)**, liver **(B)**, pancreas **(C)**, and brain **(D)**. Data are shown as mean ± standard deviation and individual animals (points). Data are shown with significance determined with a Kruskal-Wallis test with a Dunn’s test for multiple comparisons (*p<0.05, **p<0.005). Each symbol represents an individual animal.

**Supplemental Figure 4.**
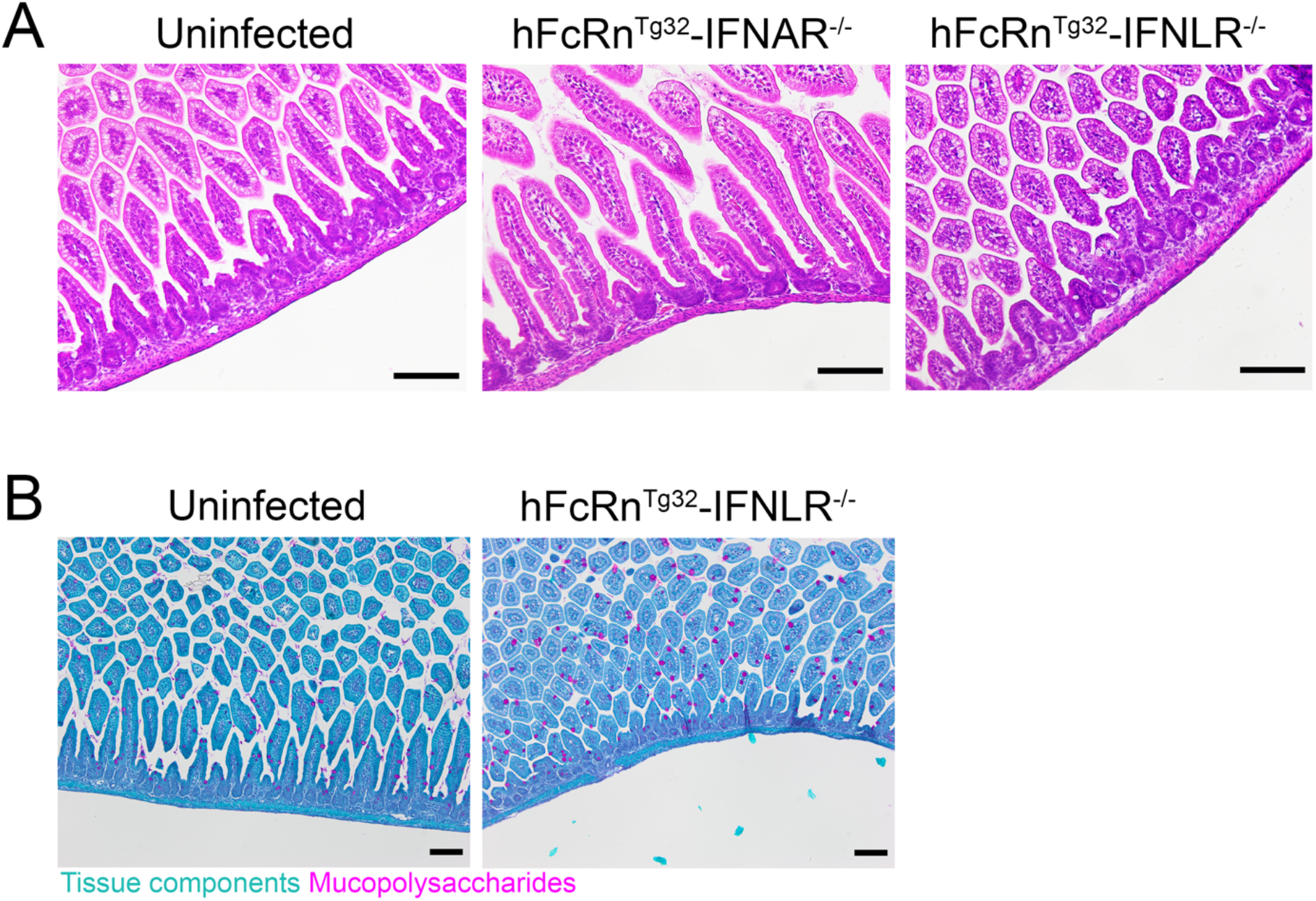
7-day old pups were orally inoculated with 10^6^ PFU of E5. At 3dpi, animals were sacrificed and intestines were collected for histology. **(A)** Hematoxylin and eosin staining of representative intestinal sections from uninfected, hFcRn^Tg32^-IFNAR^-/-^, or hFcRn^Tg32^-IFNLR^-/-^ animals. **(B)** Periodic Acid Schiff staining of representative intestinal sections uninfected or hFcRn^Tg32^-IFNLR^-/-^ animals to identify goblet cells. Scale bars at bottom right (100mm).

**Supplemental Figure 5.**
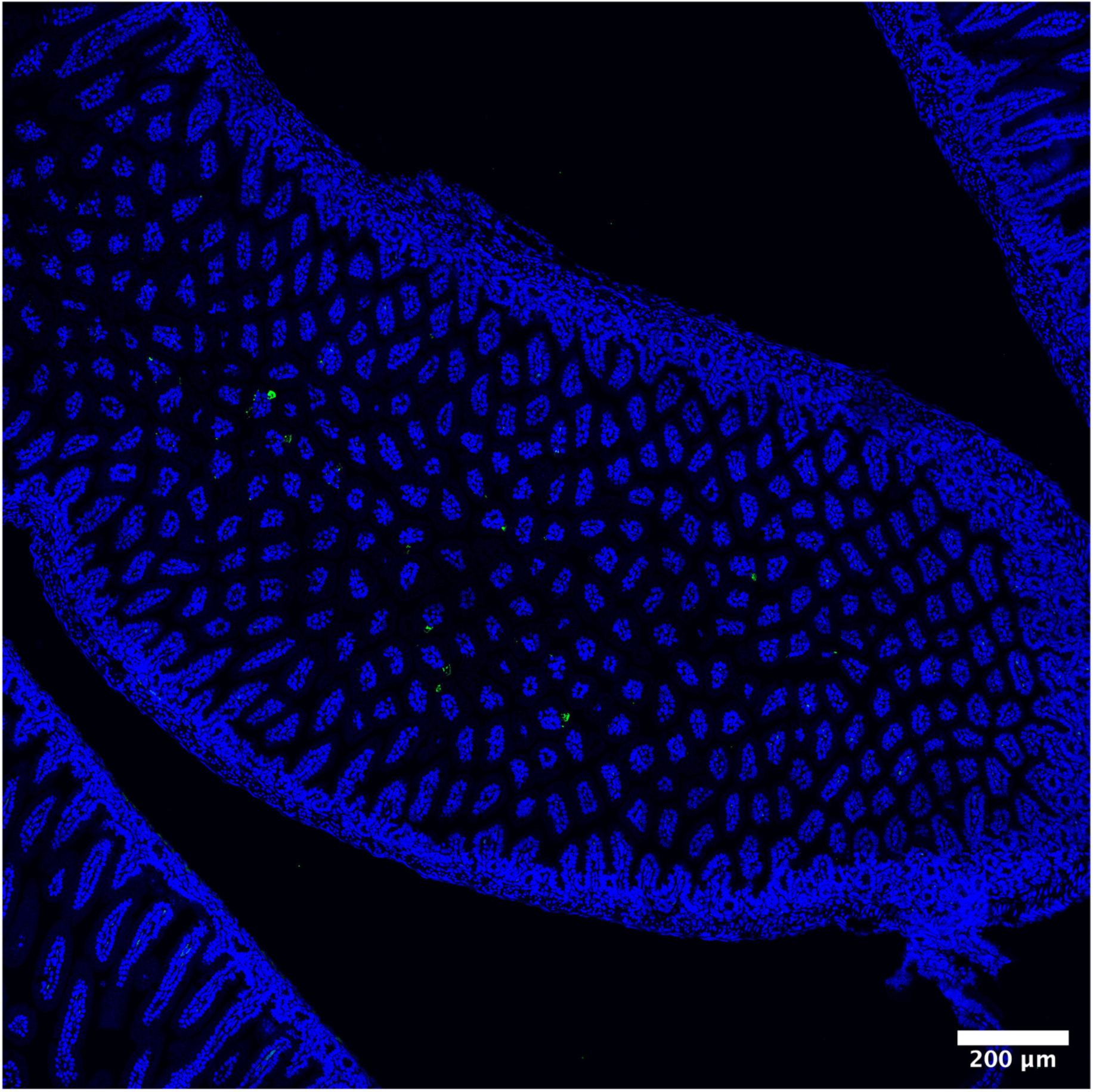
Representative tile scan of an ileum from a hFcRn^Tg32^-IFNLR^-/-^ pup with vRNA shown in green and DAPI in blue. Tile scan was done at a 20x magnification with an area of 6 by 6 tiles combined for a total of 36 individual images that were stitched together. The total area of view is 4mm^2^ per image.

## Notes

### Competing Interest Statement

The authors have declared no competing interest.

